# Decoupling of global brain activity and cerebrospinal fluid flow in Parkinson’s cognitive decline

**DOI:** 10.1101/2021.01.08.425953

**Authors:** Feng Han, Gregory L. Brown, Yalin Zhu, Aaron E. Belkin-Rosen, Mechelle M. Lewis, Guangwei Du, Yameng Gu, Paul J. Eslinger, Richard B. Mailman, Xuemei Huang, Xiao Liu

**Affiliations:** Department of Biomedical Engineering, The Pennsylvania State University, PA, USA; Department of Engineering Science and Mechanics, The Pennsylvania State University, PA, USA; Department of Neurology, Pennsylvania State University Milton S. Hershey Medical Center, Hershey, PA, USA; Department of Pharmacology, Pennsylvania State University Milton S. Hershey Medical Center, Hershey, PA, USA; Department of Radiology, Pennsylvania State University Milton S. Hershey Medical Center, Hershey, PA, USA; Department of Neurosurgery, Pennsylvania State University Milton S. Hershey Medical Center, Hershey, PA, USA; Department of Kinesiology, Pennsylvania State University Milton S. Hershey Medical Center, Hershey, PA, USA; Institute for Computational and Data Sciences, The Pennsylvania State University, PA, USA

**Keywords:** cerebrospinal fluid flow (CSF), cognitive impairment, glymphatic system, global resting-state fMRI signal, Parkinson’s disease

## Abstract

**Background:** Deposition and spreading of misfolded proteins (α-synuclein and tau) have been linked to Parkinson’s cognitive dysfunction. The glymphatic system may play an important role in the clearance of these toxic proteins via cerebrospinal fluid (CSF) flow through perivascular and interstitial spaces. Recent studies discovered that sleep-dependent global brain activity is coupled to CSF flow that may reflect glymphatic function.

**Objective:** To determine if the decoupling of brain activity-CSF flow is linked to Parkinson’s cognitive dysfunction.

**Methods:** Functional and structural MRI data, clinical motor (Unified Parkinson's Disease Rating Scale), and cognitive (Montreal Cognitive Assessment, MoCA) scores were collected from 60 Parkinson’s and 58 control subjects. Parkinson’s patients were subgrouped into those with (MoCA < 26; N = 29) and without (MoCA ≥ 26; N = 31) mild cognitive impairment (MCI).

The coupling strength between the resting-state global blood-oxygen-level-dependent signal (gBOLD) and associated CSF flow was quantified, compared among groups, and associated with clinical and structural measurements.

**Results:** gBOLD-CSF coupling decreased significantly (*p* < 0.006) in Parkinson’s patients showing MCI, compared to those without MCI and controls. Reduced gBOLD-CSF coupling was associated with decreased MoCA scores that was present in Parkinson’s patients (*p* = 0.005) but not in controls (*p* = 0.65). Weaker gBOLD-CSF coupling in Parkinson’s patients also was associated with a thinner right entorhinal cortex (Spearman’s correlation = − 0.36; *p* = 0.012), an early structural change often seen in Alzheimer’s.

**Conclusions:** The decoupling between global brain activity and associated CSF flow is related to Parkinson’s cognitive impairment.

---

Parkinson’s disease is characterized clinically by motor dysfunction ^1^ and pathologically by both α-synuclein-positive Lewy bodies/neurites and dopamine neuron loss in the substantia nigra pars compacta of the basal ganglia.^2^ Recent literature also highlights non-motor features of the disease, with cognitive decline being one of the most debilitating symptoms. Indeed, >75% of Parkinson’s patients who survive for more than 10 years ultimately develop dementia.^3^ The exact mechanisms contributing to cognitive decline in Parkinson’s disease (**PD**) are unknown and probably multifactorial. Many pathological processes have been proposed. The aggregation and spreading of α-synuclein to extra-nigral subcortical and cortical structures^4,5^ is one prominent hypothesis. Superimposed Alzheimer’s disease (AD) pathology^6–9^ could be another cause, since moderate to severe Alzheimer’s-co-pathology (e.g., amyloid-β and tau) has been found in 51% of PD patients, and is associated with faster time to dementia than Parkinson’s pathology alone.^10^

Increasing evidence suggests an important role of a newly discovered “glymphatic system” in clearing toxic proteins from the brain;^11^ these include α-synuclein,^12,13^ amyloid-β,^14^ and tau.^15^ In this clearance pathway, cerebrospinal fluid (**CSF**) flows into the interstitial space through the periarterial space, facilitated by astroglial aquaporin-4 water channels. The bulk CSF flow then drives interstitial fluid (**ISF**) flux and interstitial solutes/proteins into the perivenous space around deep-draining veins where they eventually are transported into the lymphatic system.^11,14,16^ Intriguingly, the glymphatic system functions mainly during sleep, but much less engaged during the awake state.^11^ This leads to the hypothesis that this “waste clearance” function is one universal mechanism contributing to the need for sleep in all species. This may also explain a diurnal rhythm of amyloid-β, tau, and α-synuclein.^15,17,18^

Despite its potential role in cognitive dysfunction and dementias, the study of the glymphatic system in humans has been challenging, largely due to a lack of tools that can assess glymphatic function directly and non-invasively. Two-photon microscopy and fluorescence imaging are used commonly in animal studies of the glymphatic system, but cannot yet be applied to human subjects.^19,20^ Contrast-enhanced MRI has been employed to track whole-brain CSF flow, including in human subjects,^21,22^ but broad application is limited by the need for intrathecal contrast agents and the long-duration experiments. Diffusion and structural MRI have been proposed to probe glymphatic function through surrogate markers such as perivascular space,^22–24^ however, they do not directly reflect dynamic aspects of the glymphatic system.

The pioneering study of Kiviniemi et al. ^25^ suggested that low-frequency (<0.1 Hz) resting-state fMRI (rsfMRI) blood-oxygen-level-dependent (**BOLD**) signals are linked to CSF dynamics and, thus, glymphatic function. Additional evidence supports the link between global brain activity (measured by BOLD) and glymphatic function. First, the global BOLD (**gBOLD**) signal, like glymphatic function, is highly sensitive to brain vigilance state, and also has large-amplitude fluctuations during drowsiness or sleep.^26–28^ Second, the gBOLD signal is accompanied by profound low-frequency (<0.1 Hz) physiological changes including in cardiac pulsation,^29–31^ respiration,^32–34^ and arterial signals.^35^ These latter physiological functions have been hypothesized to drive glymphatic CSF flow.^36–40^ Third, the large gBOLD signal during sleep was reported to couple to strong CSF movement^41^ that is an essential component of the glymphatic system. Collectively, the resting-state global activity and associated physiological modulations are hypothesized to represent highly coordinated neural and physiological processes closely linked to glymphatic clearance. Thus, the gBOLD-CSF coupling may serve as a marker for gauging glymphatic function.

There is limited research on the role of the glymphatic system in PD or even in PD animal models, in part because α-synuclein conformational changes and accumulation were believed to occur within neurons or at presynaptic terminals.^42^ Recent evidence, however, indicates that α-synuclein is secreted into brain extracellular spaces,^43^ suggesting a potential role of glymphatic clearance in PD. Similar to amyloid-β and tau, α-synuclein in both ISF and CSF increased in response to sleep deprivation.^15^ In addition, tau protein, which can be cleaned by glymphatic system,^44^ plays an important role in the development of PD and PD dementia.^7,45,46^ Thus, we hypothesized that coupling between the global brain signal and CSF flow is related to glymphatic function and thus cognitive impairment in Parkinson’s patients.

## Methods

### Study design and participants

Sixty PD subjects and 58 controls for whom we had rsfMRI data and Montreal Cognitive Assessment (MoCA) scores were included. The PD subjects were recruited from a tertiary movement disorders clinic and controls from the spouse population of the clinic or via IRB-approved recruitment materials posted in the local community. PD diagnosis was confirmed by a movement disorder specialist according to the UK Brain Bank criteria.^47^ All subjects were free of major/unstable medical issues such as liver, kidney, or thyroid abnormality, and deficiency of vitamin B_12_, or any cerebrovascular disease or neurological condition (other than PD). Disease duration was obtained from subject history, with onset defined as the first diagnosis by a medical professional.

### Ethics approval

The full protocol of this study was approved by the Institutional Review Board of Human Subjects Protection Office at the Penn State Milton S. Hershey Medical Center College of Medicine. All subjects provided written informed consent. The study was conducted in accordance with the principles of the Declaration of Helsinki.

### Clinical assessments

The Movement Disorder Society (MDS)-UPDRS sub-scores (UPDRS-I, UPDRS-II, and UPDRS-III) were obtained. UPDRS-I evaluates non-motor aspects of daily living, UPDRS-II assesses motor aspects of daily living reported by the subjects, and UPDRS-III evaluates motor symptoms assessed by a trained examiner.^48,49^ Levodopa-equivalent daily dosage (LEDD) was calculated for Parkinson’s subjects according to published criteria.^50^ Cognition was assessed using the Montreal Cognitive Assessment (MoCA) ^51^ and depression by the Hamilton Depression Rating Scale (HAM-D).^52^ Following procedures reported in previous studies,^51,53^ Parkinson’s patients were subgrouped into those with (MoCA < 26) and without (MoCA ≥ 26) mild cognitive impairment (MCI).

### Image acquisition and preprocessing

All MRI data were obtained on a 3.0 Tesla MR scanner (Trio, Siemens Magnetom, Erlangen, Germany, with an 8-channel phased-array head coil). To avoid systematic bias, Parkinson’s and control subjects were scanned in an intermixed fashion throughout the study. Each rsfMRI acquisition began with a T1-weighted MPRAGE structural MRI sequence (repetition time/echo time (TR/TE) = 1540/2.3 msec, field of view = 256 mm × 256 mm, matrix = 256 × 256, slice thickness = 1 mm, slice number = 176) that was used for anatomical segmentation and template normalization. In each rsfMRI session, we acquired 240 fMRI volumes using a 3D echo-planar image (EPI) sequence (flip angle = 90°, spatial resolution = 3×3×4 mm^3^, slice thickness = 3.0 mm) with TR/TE = 2000/30 msec.

RsfMRI data were preprocessed with a rsfMRI preprocessing pipeline adapted from the 1000 Functional Connectomes Project (version 1.1-beta).^54^ Specifically, we performed motion correction, skull/edge stripping, spatial smoothing (full width at maximum (FWHM) = 4mm), bandpass filtering (0.01-0.1 Hz), and linear and quadratic detrending. In addition, the first five rsfMRI volumes were discarded to avoid unsteady state of magnetization and potential edge effect of temporal filtering. These procedures generated the preprocessed rsfMRI at individual space, from which the gBOLD and CSF signals were extracted. Similar to approach of Fultz et al.,^41^ we skipped the nuisance regression of gBOLD and CSF signal since these variables were the focus of the current study. Head motion also may affect the gBOLD signal,^34^ and we thus tested its effects separately by regarding the head motion as a confounding factor as described below. To derive the fMRI-based drowsiness index ^34^ (see below), we further registered spatially the rsfMRI images to the standard MNI-152 space ^55^ (spatial resolution: 3×3×3 mm) and (temporally) standardized signals of each voxel to Z scores.

Because there is a great deal of research related to early-stage Alzheimer’s disease,^56–59^ we quantified entorhinal cortex (**ERC**) thickness and hippocampal volume from both hemispheres using Free-Surfer (version 5.1.0) from T1-weighted images.^60^ These two markers for early-stage or preclinical Alzheimer’s were selected because Parkinson’s subjects in our cohort have relatively high MoCA scores (mean MoCA was 24.8) compared to that for diagnosing Alzheimer’s (MoCA < 18).^61^ The process included motion correction, tissue classification, brain extraction, registration, and segmentation of gray matter volumetric structures, such as the ERC and hippocampus. To control for inter-subject variability of brain size, we normalized the ERC thickness and hippocampal volume with estimated total brain tissue volume.^62^

### Coupling between gBOLD signal and CSF flow

The gBOLD signal was extracted from the gray matter regions.^33,63^ It was defined based on the Harvard-Oxford cortical and subcortical structural atlases.^64^ To avoid spatial blurring from the registration process, this gray-matter mask was transformed from the MNI-152 space back to the original space of each subject based on the same rationale used in a previous study.^41^ The fMRI signal temporally was Z-normalized and then spatially averaged across all gray matter voxels. The resulting gBOLD signal reflects the amplitude and synchronization level of global brain activity. The CSF signal was extracted from the bottom slice of the fMRI acquisition near the bottom of the cerebellum and normalized by the temporal mean to percentage change. This bottom slice is expected to have maximal sensitivity to inflow effects:^65^ CSF inflow from outside the fMRI acquisition volume, which has not experienced repeated radiofrequency pulses, would increase the fMRI signal.^41^ The mask of CSF regions was obtained manually based on increased image intensity compared to surrounding brain regions in T2*-weighted fMRI and confirmed with T1-weighted MRI images.

Following procedures from Fultz et al.,^41^ the cross-correlation function first was calculated between the gBOLD and CSF signals to quantify their cross-correlations at different time lags (the gBOLD was regarded as the reference). The gBOLD-CSF cross-correlation at the time lag of +4 seconds, where the negative peak was found, was employed to quantify the gBOLD-CSF coupling for each subject. We also calculated the cross-correlation function between the negative firstorder derivative of the gBOLD signal and the CSF signal to compare with findings from Fultz et al. ^41^ We used a permutation method to test the statistical significance for the above gBOLD-CSF cross-correlations. The gBOLD and CSF signals from different subjects were matched randomly and their cross-correlation functions were computed and averaged. This procedure was repeated 10,000 times to build a null distribution for the mean gBOLD-CSF cross-correlation function, which then was used for statistical inference.

### Comparison of gBOLD-CSF coupling and association with clinical metrics

A linear regression was first conducted between the gBOLD-CSF coupling and age, and the two-sample t test then was then used to compare the gBOLD-CSF coupling between males and females. After these two initial analyses, the age- and gender-adjusted gBOLD-CSF coupling was used for all subsequent analyses described below.

We compared gBOLD-CSF coupling among control, PD-MCI (MoCA < 26), and PD-non-MCI (MoCA ≥ 26) subjects using the two-sample t-test. We then used Spearman’s correlations to quantify associations between gBOLD-CSF coupling and various clinical measures, including MoCA scores, MDS-UPDRS scores, ERC thickness, and hippocampal volumes. These correlational analyses were conducted within the control (if applicable) and PD groups, as well as the entire group of subjects.

We further evaluated the potential effects of multiple factors, including high-leverage data point, disease duration, head motion, and arousal state, on the association between the gBOLD-CSF coupling and MoCA score. We re-ran the correlation between the two after excluding two high-leverage data points with MoCA less than 15, as well as controlling for disease duration. We estimated head motion and drowsiness/sleepiness level of subjects and then re-tested the association between gBOLD-CSF coupling and MoCA controlling for these factors. For head motion, we quantified framewise displacement (FD), is defined as the sum of the absolute values of six translational and rotational motion parameters.^66^ These values then were averaged within the same session to quantify individual subject head motion.^34^ For the arousal state, we adapted an fMRI-based drowsiness index.^34,67^ Briefly, we computed the spatial correlation between each rsfMRI volume and a pre-defined drowsiness template derived previously ^28^ and then took the envelope amplitude of this spatial correlation (i.e., similarity) as an estimation of instantaneous brain arousal level. This drowsiness index then was averaged within each session to represent the overall drowsiness level of each subject and included in the correlation analysis.

### Statistical analysis

In summary, the two-sample t-test was used for group comparisons of continuous demographic and clinical measures, whereas the Fisher exact test was utilized for categorical metrics.^68^ The permutation test was used to assess the statistical significance of the gBOLD-CSF correlations tested. Spearman’s correlation was used to quantify inter-subject associations between the gBOLD-CSF coupling and various clinical measures in order to account for potential non-normality of these variables. P-values < 0.05 were considered statistically significant. We also corrected multiple comparisons using the Bonferroni method.^69^

## Results

### Participant characteristics

Demographic and clinical data are presented in **Table 1**. PD participants on average were ca. 3-years older than controls (*p* = 0.027). The two groups had similar numbers of males and females (*p* = 0.58). PD participants had an average disease duration of 6.6 ± 5.7 years, and significantly higher HAM-D (*p* = 4.3×10^−5^) and lower MoCA scores (*p* = 0.009) than controls. There were approximately equal numbers of normal cognition PD patients (MoCA ≥ 26, N = 31) and those with MCI (MoCA < 26, N = 29).

**Table 1:** Characteristics of the cohort.

|  | **Controls (N=58)** | **PD (N=60)** | **P value** |
| --- | --- | --- | --- |
| Age, y | 61.4 (8.0) | 64.7 (8.2) | 0.027 |
| Gender (Male/ Female) | 31/27 | 6/24 | 0.58 |
| Disease duration, y | — | 6.6 (5.7) | — |
| UPDRS-I | — | 10.6 (7.9) | — |
| UPDRS-II | — | 9.4 (8.5) | — |
| UPDRS-III | — | 22.2 (20.0) | — |
| LEDD, mg | — | 736 (487) | — |
| MoCA | 26.5 (2.6) | 24.8 (4.1) | 0.009 |
| HAM-D | 4.0 (3.3) | 7.8(4.6) | $4.3 \times 10^{-5}$ |

*Data represent the mean and standard deviation (in parentheses) unless otherwise indicated.*
*Pairwise comparisons were performed based on the two-sample t-test for all measures except for* *gender that used the Fisher exact test.*
*Abbreviation: PD, Parkinson's disease group; UPDRS: Unified Parkinson's Disease Rating* *Scale; LEDD: levodopa equivalent daily dose; MoCA: Montreal Cognitive Assessment; HAM-D:* *Hamilton Depression Rating Scale.*

### gBOLD signal is coupled to CSF signal changes

We first examined whether the gBOLD signal is coupled to strong CSF signal changes in this Parkinson’s dataset. As shown in a representative control, changes in the gBOLD signal are associated with large CSF signal changes (**Fig. 1B**, black and gray arrows). The mean gBOLD-CSF cross-correlation function was characterized by a positive peak (*r* = 0.28; *p* < 0.0001; permutation test, *N =* 10,000) around the −4-sec lag (i.e., correlation with shifting CSF ahead of time by ca. 4 sec), whereas there was a negative peak (*r* = −0.28; *p* < 0.0001) at the lag of +4 sec (**Fig. 1C**, left panel). The cross-correlation function between the CSF signal and the negative first-order derivative of the gBOLD signal displayed a large positive peak at the zero time-lag (*r* = 0.37; *p* < 0.0001). Overall, both cross-correlation functions (**Fig. 1C**) showed patterns consistent with the data of Fultz et al.,^41^ confirming a systematic coupling between the global brain signal and CSF flow.

**Fig. 1.**
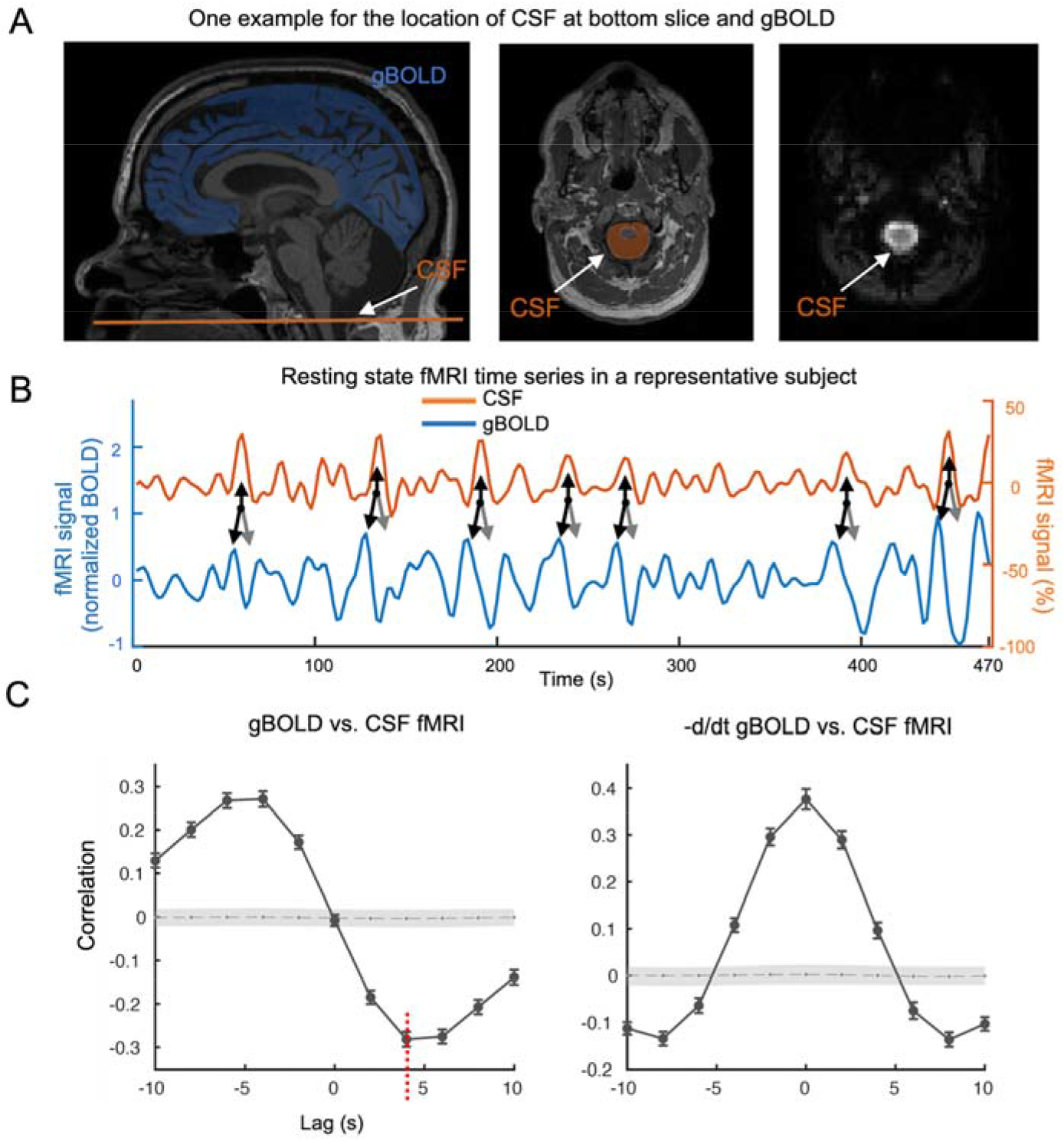
Systematic coupling between the gBOLD and CSF signals measured by rsfMRI. (**A**) The gBOLD and CSF fMRI signals were extracted from the whole-brain gray matter regions (the blue mask on a T1-weighted structural MRI in the left panel) and the CSF region at the bottom slice of the fMRI acquisition (the bright region in the right panel and the orange-shaded area in the middle panel), respectively. (**B**) The gBOLD and CSF signal from a representative subject (control) showed corresponding changes of large amplitude. Large CSF peaks (upwards black arrows) often are preceded by a large positive gBOLD peak (downwards black arrows) and followed by a large negative gBOLD peak (gray arrows). (**C**) The averaged cross-correlation function (N =118 subjects) between the gBOLD signal (reference) and the CSF signal (left), a well as between the first-order negative derivative of gBOLD signal (reference) and the CSF signal (right). The gray dashed line and the shaded region mark 95% confidence intervals for the mean correlation computed on shuffled signals (see **Materials and methods** for details). The cross-correlations between the gBOLD and CSF signal at the +4 seconds lag (mean *r*: −0.28; *p* < 0.0001, permutation test with N =10,000; red dashed line) was used to represent “the gBOLD-CSF coupling” for subsequent analyses.

### Comparison of gBOLD-CSF coupling among groups

We tested whether the gBOLD-CSF coupling was affected by age or gender. A significant correlation was found between age and gBOLD-CSF coupling (**Fig. 2A**; ρ = 0.32, *p* < 0.001), with increased age associated with weaker (less negative) gBOLD-CSF coupling. No significant difference (*p* = 0.44) in gBOLD-CSF coupling was found between males and females. For scientific rigor, we controlled for the age and gender effects on gBOLD-CSF coupling for all subsequent analyses.

**Fig. 2.**
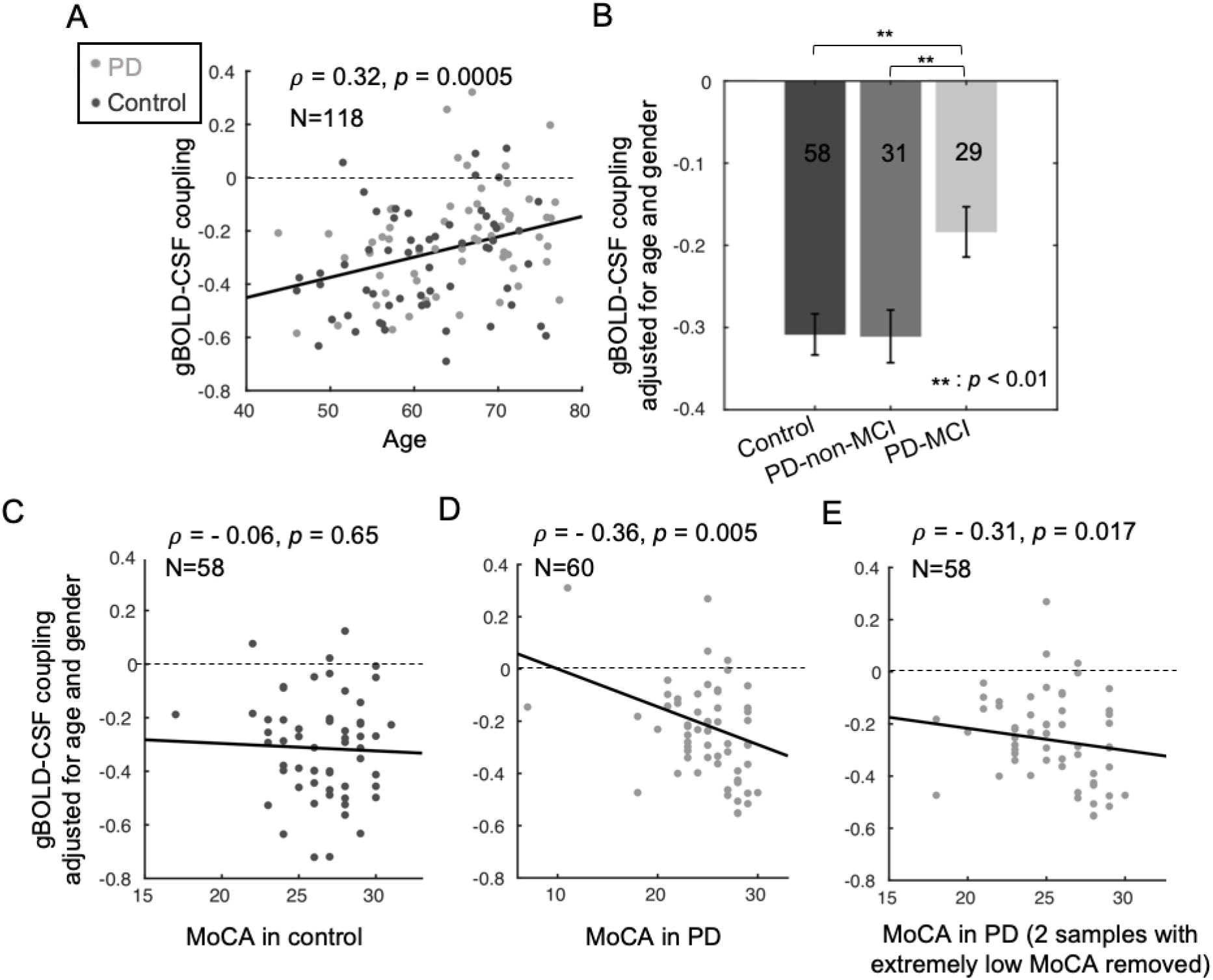
The associations of gBOLD-CSF coupling to age, disease condition, and MoCA. (**A**) Older subjects have a weaker (less negative) gBOLD-CSF coupling (Spearman’s ρ = 0.32, *p* = 0.0005). (**B**) Compared with controls and PD-non-MCI subjects, PD-MCI subjects have decreased gBOLD-CSF coupling strength (adjusted for age and gender; *p* ≤ 0.006, two-sample t-test). (**C-D**) The correlation between the gBOLD-CSF coupling (adjusted for age and gender) and MoCA scores is significant in the PD group (ρ = −0.36, *p* = 0.005, *p_Bonferroni_* = 0.01; **D**) but not in the controls (ρ = −0.06, *p* = 0.65; **C**). (**E**) The significant correlation between gBOLD-CSF coupling and MoCA within PD subjects remained significant when removing the two subjects with extremely low MoCA scores (ρ = −0.31, *p* = 0.017, *p_Bonferroni_* = 0.034). The dashed lines indicate where the gBOLD-CSF coupling equals zero.

The gBOLD-CSF coupling metric then was compared among groups. PD-MCI subjects showed significantly weaker gBOLD-CSF coupling (**Fig. 2B**) than controls (*p* = 0.003) and PD-non-MCI (*p* = 0.006) subjects. gBOLD-CSF coupling strength, however, was not different between control and PD-non-MCI subjects (*p* = 0.96).

### Association between gBOLD-CSF coupling and cognitive and PD measures

The gBOLD-CSF coupling then was correlated with various clinical and structural measurements. Consistent with comparisons across disease groups, weaker (less negative) gBOLD-CSF coupling was correlated significantly with lower MoCA scores across all PD subjects (ρ = −0.36, *p* = 0.005, *p_Bonferroni_* = 0.01, **Fig. 2D**) but not across the controls (ρ = −0.06, *p* = 0.65, **Fig. 2C**). The Parkinson’s-specific gBOLD-CSF coupling/MoCA associations remained significant (ρ = −0.31, *p* = 0.017, *p_Bonferroni_* = 0.034, **Fig. 2E**) when excluding two subjects with very low MoCA scores. The association is also significant (*p* < 0.05) across the entire group of subjects (**Suppl. Fig. 1**), or after controlling for disease duration (**Suppl. Fig. 2**), head motion (**Suppl. Fig. 3**), and drowsiness score (**Suppl. Fig. 4**). As seen **Suppl. Fig. 5**, there was no significant correlation found between the gBOLD-CSF coupling and any of the MDS-UPDRS motor (UPDRS-II, *p* = 0.80, UPDRS-III, *p* = 0.27); or non-motor subscores UPDRS-I, *p* = 0.49).

### Association of BOLD-CSF coupling strength with structural changes in Parkinson’s

Weaker (less negative) gBOLD-CSF coupling was associated significantly with a thinner right ERC region (ρ = −0.36, *p* = 0.012, *p_Bonferroni_* = 0.024, **Fig. 3A**), whereas only a trend was found for the left ERC thickness (ρ = −0.19, *p* = 0.19, *p_Bonferroni_* = 0.38, **Fig. 3B**). Weaker gBOLD-CSF coupling was not associated with lower hippocampal volume bilaterally (*p* > 0.05; **Suppl. Figs. 6A-B**). There was no correlation between gBOLD-CSF coupling and ERC thickness or hippocampal volume in the control group (*p* > 0.05; see **Suppl. Fig. 7**).

**Fig. 3.**
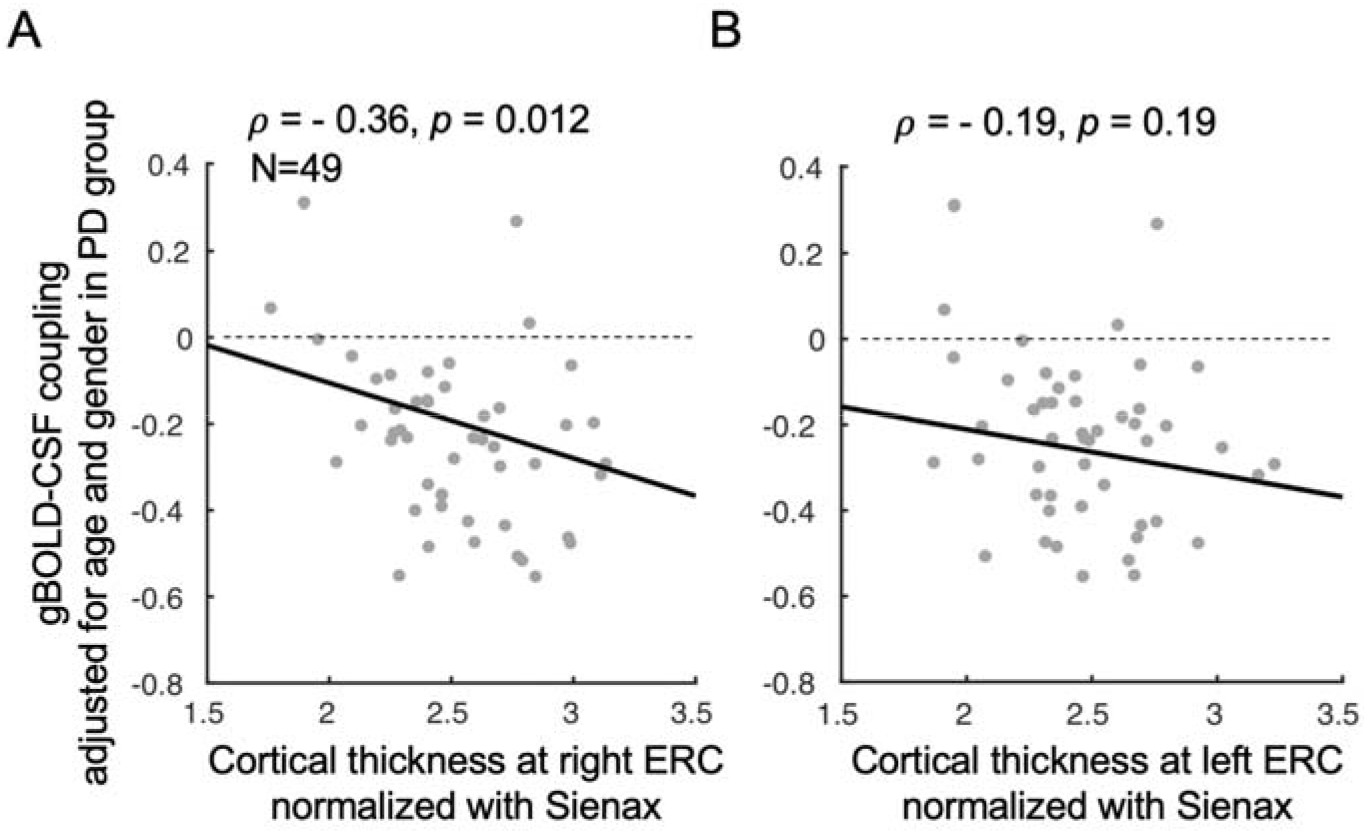
Associations between gBOLD-CSF coupling and thickness of the entorhinal cortex in PD patients. The PD patients with thinner right ERC tend to have weaker gBOLD-CSF coupling (Spearman’s ρ = −0.36, *p* = 0.012, *p_Bonferroni_* = 0.024) (**A**). This association is similar but not significant (ρ = −0.19, *p* = 0.19, *p_Bonferroni_* = 0.38) for the left ERC (**B**). The ERC thickness was normalized with the Sienax index.^62^

## Discussion

The current study demonstrated a strong coupling between global brain activity and CSF flow using rsfMRI data obtained both from Parkinson’s and control subjects, consistent with the idea that gBOLD-CSF coupling is related to glymphatic function. The strength of this gBOLD-CSF coupling was significantly lower in those Parkinson’s patients with cognitive impairment compared to either controls or Parkinson’s patients without cognitive dysfunction. Moreover, the gBOLD-CSF coupling strength was correlated with cognitive scores, but only in Parkinson’s patients and not controls, suggesting it is linked to Parkinson’s-related processes. No correlation was found between gBOLD-CSF coupling and UPDRS motor or non-motor scores, suggesting it is related specifically to a cognitive component of Parkinson’s. The reduced gBOLD-CSF coupling also was associated with a thinner ERC that has been observed in early-stage Alzheimer’s, suggesting a possible contribution from Alzheimer’s-like pathology in PD-MCI. These findings suggest that gBOLD-CSF coupling is a potential noninvasive marker for the glymphatic function that may be related to dementia pathology.

### gBOLD-CSF coupling may be related to the sleep-dependent glymphatic system

Resting-state gBOLD-CSF coupling likely results from highly coordinated neural and physiological processes that are linked closely to glymphatic clearance. This potential relationship is supported by at least three lines of evidence. First, the resting-state gBOLD signal shows a similar sleep dependency as glymphatic function, and is strongly promoted by drowsiness and sleep.^26,27,70–72^ Sleep deprivation ^71^ and hypnotic drugs (e.g., midazolam),^73^ significantly enhance the gBOLD signal, whereas caffeine has the opposite effect.^72^ It became clear recently that the strong gBOLD signal fluctuation reflects transient arousal modulations of 10-20 sec.^28^ Second, this gBOLD signal is accompanied by slow (<0.1 Hz), but strong physiological modulations of cardiac,^29–31^ respiratory,^33,34^ and arterial^35^ signals. These physiological modulations have been hypothesized to be the main drivers of glymphatic CSF flow.^36–40^ It is likely that they are mediated through the autonomic system via transient arousal variations underlying the gBOLD signal.^29^ In at least one mouse model (for cerebral amyloid angiopathy), the low-frequency modulations in vessel tone has been linked to amyloid-β clearance.^74^ Thus, the gBOLD signal and associated physiological modulations should provide sleep-dependent dynamics for CSF flow that are essential for the glymphatic system. Indeed, we found that strong CSF movements coupled to the large gBOLD signal,^41^ adding a new piece of evidence for their relevance to the glymphatic system. This coupling relationship was confirmed and extended in the present study of Parkinson’s patients. Overall, this potential link to the glymphatic system could be the primary explanation for the association we found between gBOLD-CSF coupling and cognitive dysfunction in Parkinson’s.

### gBOLD-CSF coupling may be a new functional metric for cognitive decline

The introduction summarized evidence for the glymphatic system being involved in clearance of toxic misfolded proteins such as α-synuclein, amyloid-β and tau, thereby possibly contribute to the cognitive decline in PD patients. The initial test of this hypothesis in the current study indicated that gBOLD-CSF coupling was decreased significantly only in Parkinson’s patients with cognitive decline. It is also correlated with cognitive MoCA scores, but not with motor and other non-motor symptoms measured by MDS-UPDRS scores. Together, the data provide the first evidence that gBOLD-CSF coupling may serve as a new, non-invasive, functional metric for quantifying cognitive decline, and warrants future studies of its relationship to the ultimate development of dementia in PD.

### gBOLD-CSF coupling may reflect aspects of Alzheimer’s pathology

The present study showed that weaker gBOLD-CSF coupling was associated with a thinner right ERC, although correlations with left ERC thickness or bilateral hippocampal volumes did not reach statistical significance. Right ERC thinning and early loss of ERC symmetry has been suggested a marker for preclinical Alzheimer’s.^75,76^ A more pronounced loss in right ERC thickness also is seen with aging.^77^ In Alzheimer’s patients, gray matter loss in the right ERC has been shown to occur earlier and be larger than in other brain regions.^75^ In fact, the ERC is known to be the earliest affected brain region in both Alzheimer’s and Parkinson’s.^78–80^ Thus, decreased gBOLD-CSF coupling may be an indicator of early cognitive decrements, and its correlation with cognitive impairment in PD-MCI may be attributed partly to structural changes similar to those in in early Alzheimer’s.

### Limitations and future directions

The present study linked the gBOLD-CSF coupling to cognitive dysfunction in PD, but one clear limitation of the present study was that it did not directly link the gBOLD-CSF coupling to accumulation of toxic chemical species (proteins or small molecules). This provides a fertile area for mechanistic understanding of the association of gBOLD-CSF coupling with cognitive impairment. Future studies should also explore other imaging markers that result from the same low-frequency global brain activity and physiological modulations, and relate them to cognitive decline and dementias. Related experiments could include quantification of α-synuclein and tau (either with PET imaging or CSF measurements), metabolomic studies of CSF, and longitudinal studies of how gBOLD-CSF coupling relates to the prognosis and progression of Parkinson’s disease dementia. Finally, despite the recent evidence, the relationship between gBOLD-CSF coupling and the glymphatic system remains hypothetical, and direct evidence using both murine and non-human primate models will be very important.

## Acknowledgements

We thank Jiaxuan Diao for preparing some figures and reviewing the manuscript.

## Author Roles

Research project: A. Conception, B. Organization, C. Execution;

Statistical Analysis: A. Design, B. Execution, C. Review and Critique;

Manuscript Preparation: A. Writing of the first draft, B. Review and Critique;

**FH**: 1A, 1B, 1C, 2B, 2C, 3A;

**GLB**: 1C, 2C, 3A, 3B;

**YZ**: 1C, 2B;

**AEBR**: 1C, 2B;

**MML**: 2C, 3A, 3B;

**GD**: 2C, 3B;

**YG**: 1C, 2C;

**PJR**: 3B;

**RBM**: 2C, 3B;

**XH**: 1A, 1B, 2A, 2C, 3B;

**XL**: 1A, 1B, 2A, 2C, 3A, 3B.

## Financial disclosure and conflict of interest statement

This work was supported by the National Institutes of Health (NIH) Pathway to Independence Award (K99/R00 5R00NS092996-03), the Brain Initiative award (1RF1MH123247-01), and the NIH R01 award (1R01NS113889-01A1). This work was also supported by the National Institute of Neurological Disorders and Stroke Parkinson’s Disease Biomarker Program (NS06722 and NS112008 to X.H), the National Institute of Aging (AG067651 to G.B), the Hershey Medical Center General Clinical Research Center (National Center for Research Resources, Grant UL1 RR033184 that is now at the National Center for Advancing Translational Sciences, Grant UL1 TR000127), the PA Department of Health Tobacco CURE Funds, and the Penn State Translational Brain Research Center. The authors report no financial interests or potential conflicts of interest.

## Supplemental files

**Suppl. Fig. 1.**
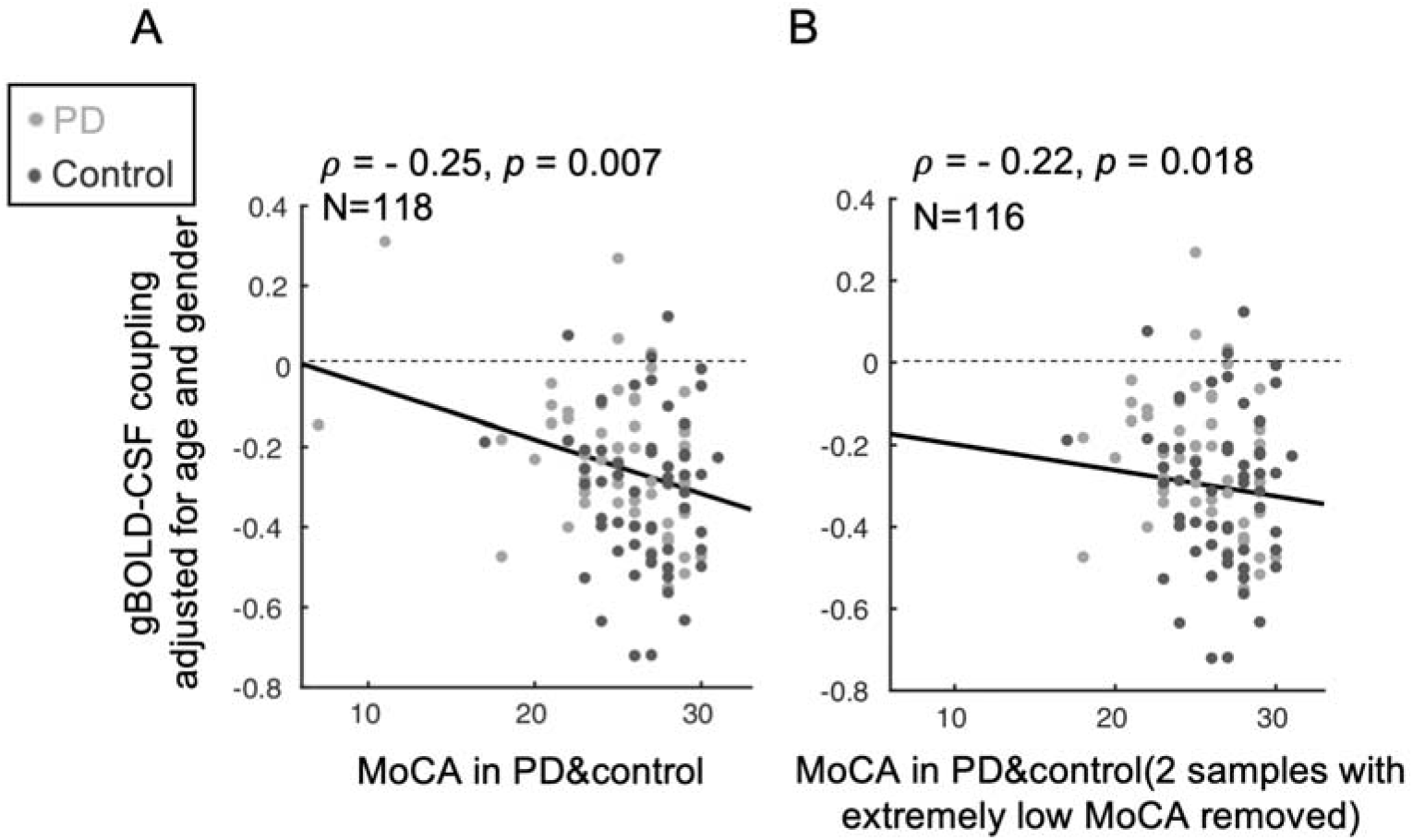
The association between the gBOLD-CSF coupling and MoCA remains significant (*p* < 0.018) in the entire cohort of subjects (**A**) and after removing two high leverage samples (from two PD subjects) with extremely low MoCA scores (**B**).

**Suppl. Fig. 2.**
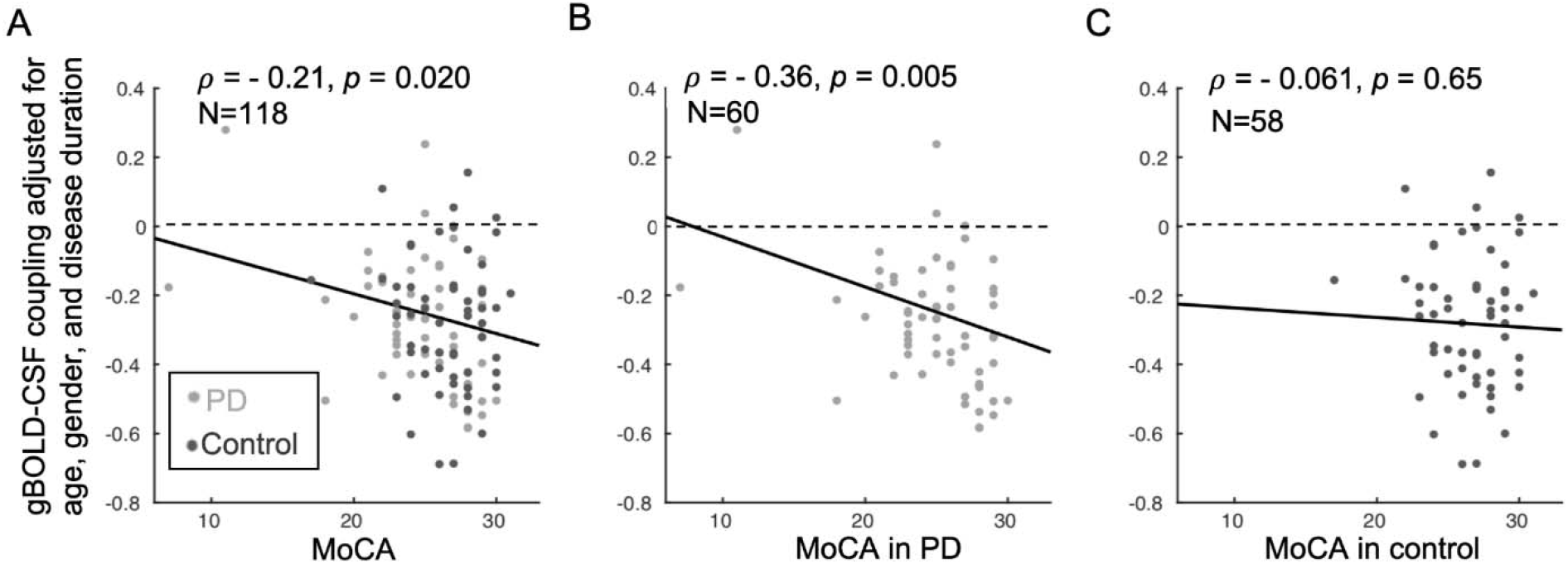
The associations between the gBOLD-CSF coupling and MoCA remains significant (*p* < 0.05) for both the entire group of subjects (**A**) and the PD group (**B**), after controlling the age, gender, and disease duration, while this significant correlation cannot be found in controls (**C**).

**Suppl. Fig. 3.**
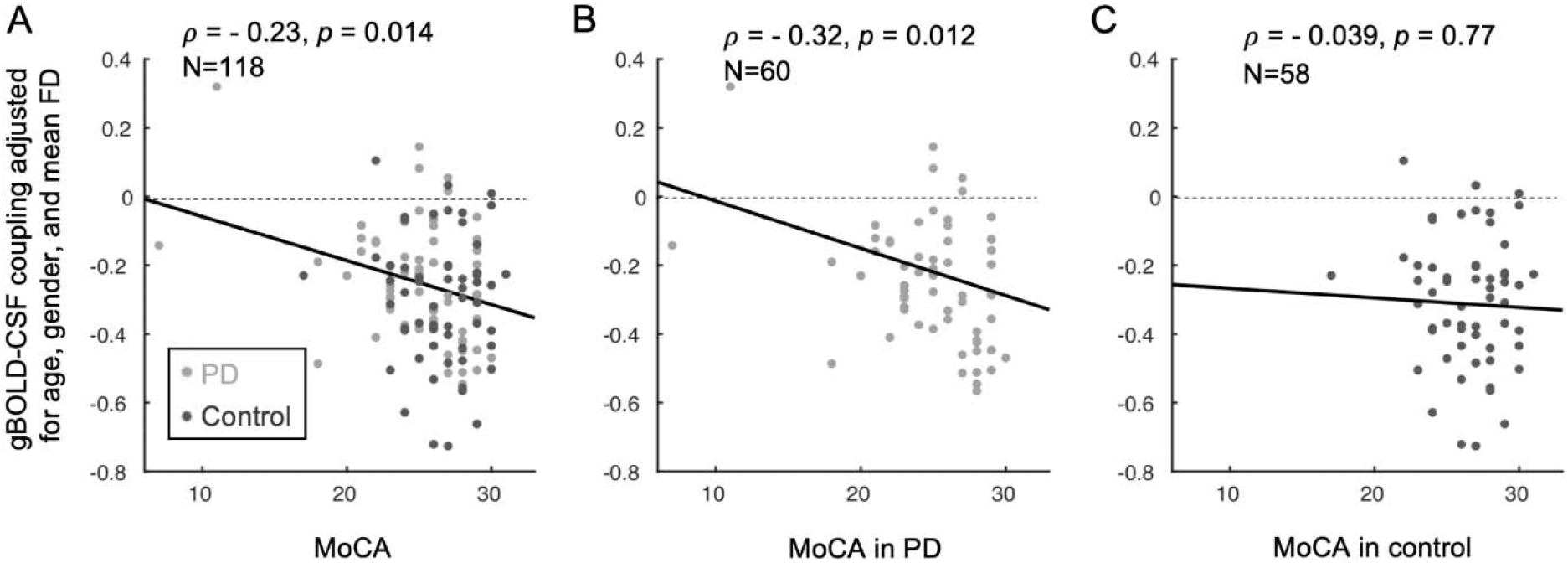
The association between the gBOLD-CSF coupling and MoCA remains significant (*p* < 0.05, corrected) for both the overall group of subjects (**A**) and within Parkinson’s subjects (**B**), after controlling the age, gender, and head motion, whereas the correlation was absent in controls (**C**).

**Suppl. Fig. 4.**
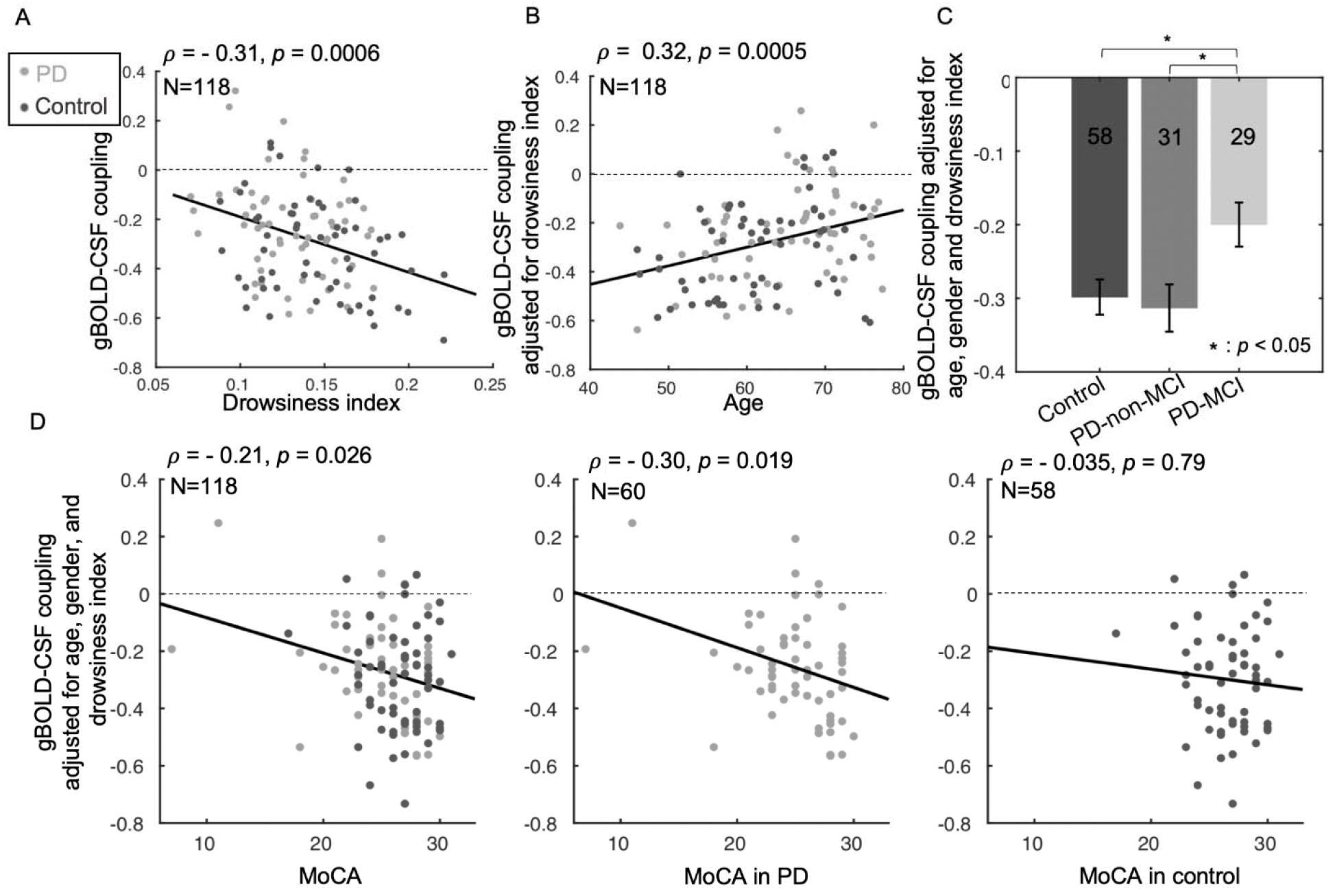
The association between the sleep-dependent gBOLD-CSF coupling and age, group, and MoCA scores is not driven by changes in the drowsiness index. (**A**) The gBOLD-CSF coupling was correlated significantly with the drowsiness index (increased drowsines index corresponds to a drowsier state) in all subjects, which is quantified based on the similarity between the rsfMRI time-series and a drowsiness template. (**B**) When adjusted for the drowsiness index, weaker gBOLD-CSF coupling strength remained significantly associated with increasing age (ρ = 0.32, *p* < 0.001). (**C**) PD-MCI subjects had decreased gBOLD-CSF coupling compared to control or PD-non-MCI after adjusting for the drowsiness index (both *p* ≤ 0.014). (**D**) Similar to **Fig. 2C**, gBOLD-CSF coupling (age, gender, and drowsiness index adjusted) remained significantly correlated with MoCA scores across both the entire group of subjects and within Parkinson’s but not for controls. The dashed lines indicate where the gBOLD-CSF coupling equals zero.

**Suppl. Fig. 5.**
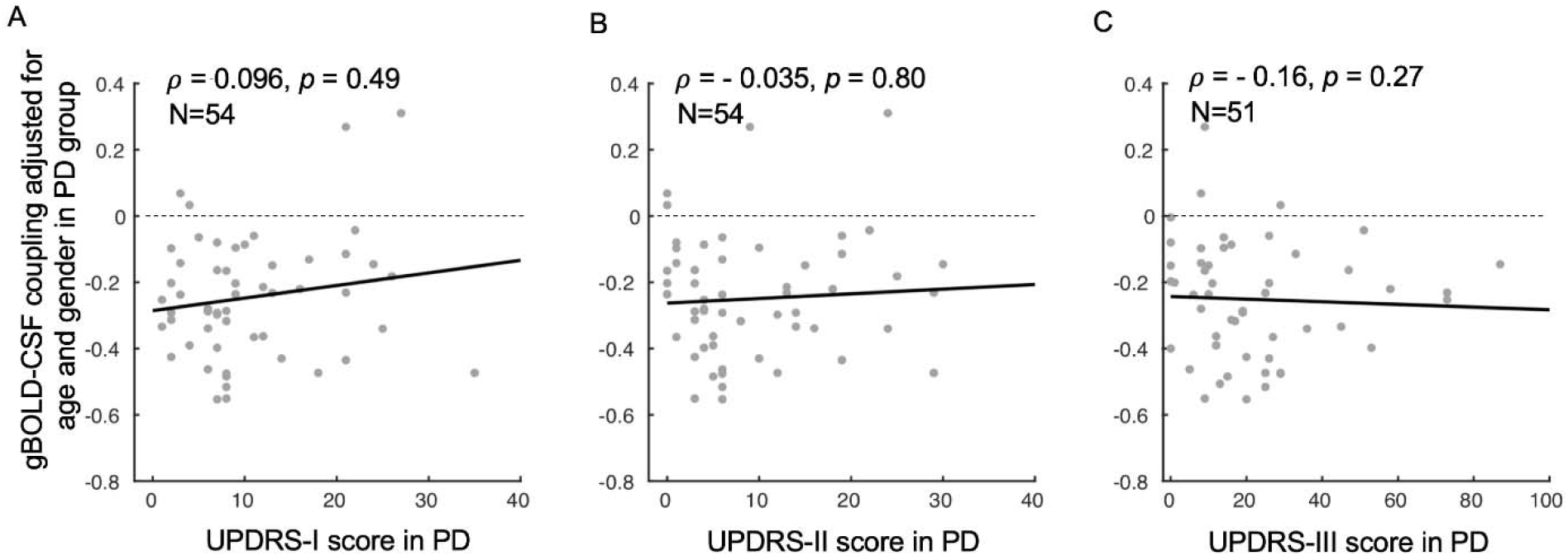
The associations (linear regression) between the gBOLD-CSF coupling (adjusted for age and gender) and MDS-UPDRS scores were not significant (*p* > 0.05) within Parkinson’s subjects. The linear regression lines were estimated based on least-squares fitting.

**Suppl. Fig. 6.**
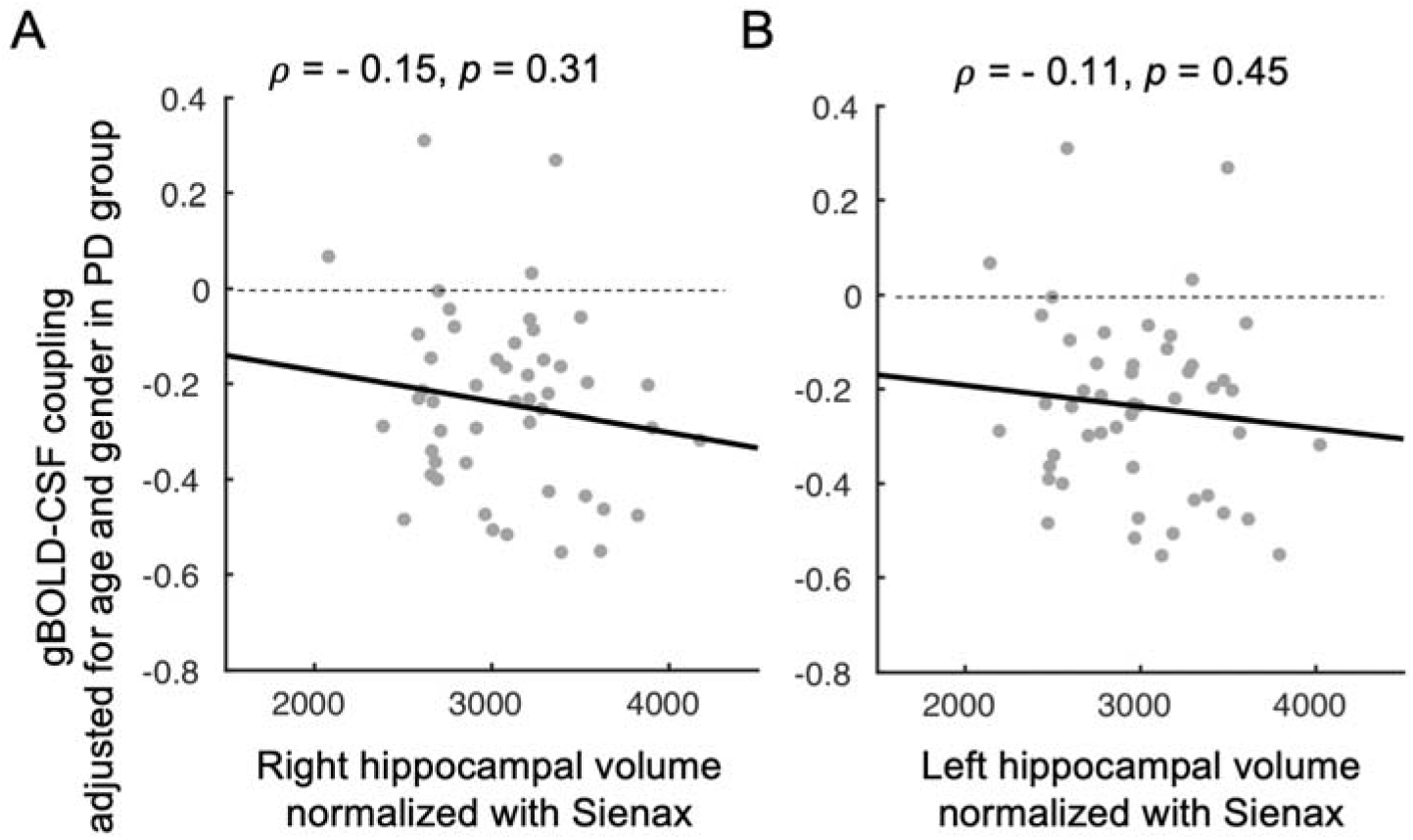
Associations between gBOLD-CSF coupling and structural measures of the hippocampus in PD patients. (**A**-**B**) The gBOLD-CSF coupling showed a negative correlation with the bilateral hippocampal volumes (normalized with the Sienax index),^62^ but none of the correlations were statistically significant (*p* > 0.05).

**Suppl. Fig. 7.**
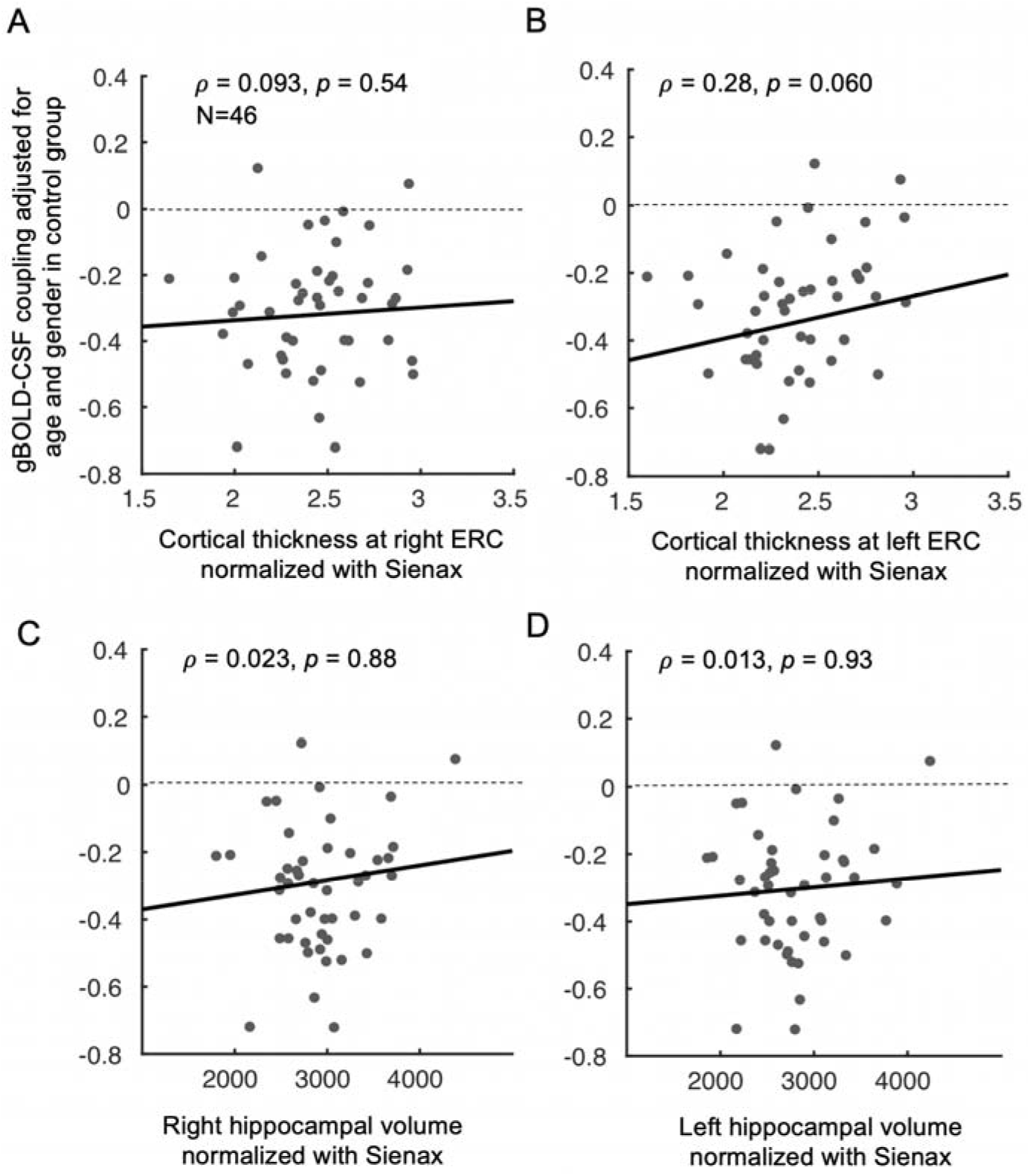
gBOLD-CSF coupling and structural measures in the entorhinal cortex and hippocampus were not correlated in controls (all *p* > 0.05). The linear regression lines were estimated based on least-squares fitting.

## Notes

### Competing Interest Statement

The authors have declared no competing interest.

